# Functional characterization of mutants deficient in N-terminal phosphorylation of the chloroplast ATP synthase subunit β

**DOI:** 10.1101/2021.11.04.467332

**Authors:** Deserah D. Strand, Daniel Karcher, Stephanie Ruf, Anne Schadach, Mark A. Schöttler, Omar A. Sandoval Ibanez, Ralph Bock

**Author notes:** DOE Plant Research Laboratory, 612 Wilson Rd 106, Michigan State University, East Lansing, MI, 48824, USA. CAuthor for contact.

## Abstract

Understanding the regulation of photosynthetic light harvesting and electron transfer is of great importance to efforts to improve the ability of the electron transport chain to supply downstream metabolism. The central regulator of the electron transport chain is the ATP synthase, the molecular motor that harnesses the chemiosmotic potential generated from proton coupled electron transport to synthesize ATP. The ATP synthase is regulated both thermodynamically and post-translationally, with proposed phosphorylation sites on multiple subunits. In this study we focused on two N-terminal serines on the catalytic subunit β, previously proposed to be important for dark inactivation of the complex to avoid ATP hydrolysis at night. Here we show that there is no clear role for phosphorylation in the dark inactivation of ATP synthase. Instead, mutation of one of the two phosphorylated serine residues to aspartate strongly decreased ATP synthase abundance. We propose that the loss of N-terminal phosphorylation of ATPβ may be involved in proper ATP synthase accumulation during complex assembly.

## Introduction

Photosynthesis provides the starting material for nearly all life on earth. There is currently a drive to improve photosynthesis to meet food demands of a growing global population and make renewable biofuels to combat climate change. Understanding photosynthesis and how it is regulated is important if we want to modify these reactions to increase plant growth or increase flexibility to respond to stress conditions (Ort et al., 2015).

The so-called ‘light reactions’ of photosynthesis are comprised of the thylakoid embedded electron transport chain that feeds the Calvin-Benson-Bassham cycle with the ATP and NADPH required for CO_2_ fixation. The central regulator of the thylakoid reactions of photosynthesis is the ATP synthase (discussed in (Kramer et al., 2004)). ATP synthase is a molecular motor comprised of a membrane associate CF_o_ subcomplex and a soluble CF_1_ subcomplex that work together to convert protonmotive force (*pmf*), generated by proton coupled electron transfer, into phosphorylation potential in the form of ATP (Hahn et al., 2018).The ΔpH component of *pmf* activates pH-dependent feedback regulation of light harvesting (i.e., exciton quenching, q_E_), the largest component of non-photochemical quenching (NPQ) and lowers the rate constant for deprotonation of plastoquinol at the cytochrome *b_6_f* (*bf*) complex Q_o_ site (photosynthetic control) (Eberhard et al., 2008). In high light, q_E_ and photosynthetic control protect photosystem II and I, respectively, from photodamage (Li et al., 2002; Suorsa et al., 2012). Changing the activation state of the ATP synthase has a rapid direct effect on these photoprotective processes (Kanazawa and Kramer, 2002; Takizawa et al., 2008; Zhang et al., 2009), and the understanding of ATP synthase regulation is important for our understanding of the regulation of photosynthesis as a whole.

While regulation of ATP synthase is still an open topic for investigation, it is likely that under standard growth conditions gene expression has little to no role in the control of ATP synthesis kinetics in mature leaves (Rott et al., 2011). Instead, ATP synthase is regulated by substrate availability, protein-protein interactions, and multiple post-translational modifications. While it is unlikely that, in the light, thermodynamics would favor ATP hydrolysis, metabolic changes may still contribute to altered kinetics under conditions where substrate is limited (Kanazawa and Kramer, 2002; Takizawa et al., 2008). In the dark, the ATPg subunit (encoded by the *ATPC1* gene in the nuclear genome) is oxidized, and leads to a decrease in the rate constant of proton efflux (Wu et al., 2007; Kohzuma et al., 2013).

An additional post-translational modification proposed to modulate ATP synthase activity during light to dark transitions and substrate limitation is phosphorylation (Bunney et al., 2001; Takizawa et al., 2008; Reiland et al., 2009). While there have been multiple proposed phosphorylation sites on ATP synthase, two serine residues in the N-terminal domain of the chloroplast-encoded CF_1_-β subunit (encoded by *atpB*) have raised particular interest, because they were shown to be phosphorylated at the end of the night (del Riego et al., 2006; Reiland et al., 2009). Due to the timing of the phosphorylation, it was proposed that these residues are specifically involved in the inactivation of ATP synthase in the dark to avoid ATP hydrolysis. In this work, we set out to test this hypothesis by mutating these two residues (serine 8 and serine 13) to either an alanine, to eliminate the phosphorylation site, or an aspartic acid, to mimic constitutive phosphorylation. To assess the function of each of these residues in ATP synthase dark inactivation, we used chloroplast transformation to generate a set of mutants with all possible combinations of phosphorylation states

## Results

### Generation of transplastomic tobacco plants with mutated phosphorylation sites in the atpB gene

To test the hypothesis that phosphorylation of the N-terminus of ATPβ is involved in regulation of ATP synthase, we introduced mutations either abolishing the phosphorylation site (S-to-A) or mimicking a phosphorylated residue (S-to-D). To achieve this, we used co-transformation of 9 plasmids: 8 plasmids containing our mutations of interest in all combinations, and one plasmid with the *aadA* selectable marker gene cassette. This allowed us to identify plants resistant to spectinomycin (conferred by the *aadA* gene; (Svab and Maliga, 1993; Bock, 2015)), and then further screen these transplastomic plants for the presence of any of the desired mutations, allowing for isolation of multiple mutations within the same transformation experiment. After biolistic bombardment and selection, the first round of plant regenerants were screened by PCR and sequencing of the *atpB* gene (encoding ATPβ). Plants that contained the desired mutation were sub-cultured further, and the second round of regenerated plants were screened again by PCR and DNA sequencing. Plants appearing to have a single peak in the Sanger sequencing chromatogram were further analyzed by Southern blotting for homoplasmic incorporation of the selection cassette. Plants that appeared to be homoplasmic for both the *atpB* mutation and the co-transformed selection cassette were transferred to the greenhouse for seed production.

To determine homoplasmy, seeds from these plants were plated on spectinomycin and streptomycin and considered homoplasmic for the selection cassette when all seedlings were indistinguishable from the wild type (WT) grown on non-selective medium (Bock, 2001). To ultimately confirm homoplasmy and stability of the introduced mutations, ~20 of these seedlings were pooled and sequenced for the desired mutations in *atpB* via PCR amplification and Sanger sequencing. Plant lines with single peaks at the site(s) of the desired mutation(s) were considered homoplasmic (Figure 1A and Supplemental Figure 1). DNA from pooled seedlings was also used for final Southern blot analysis to demonstrate the homoplasmic presence of the selectable marker gene *aadA* in the plastid genomes by a 1.2 kb increase in band size using a probe against the *psaB* gene (Figure 1B). At least three independently transformed transplastomic lines were isolated for each mutation. To assay the effect of the mutations on transcript abundance, we performed a northern blot for the *atpB* transcript with total RNA isolated from young leaves. This analysis revealed that there is no apparent reduction in transcript abundance (Figure 1C), indicating there are no apparent changes in transcription or transcript stability due to a specific mutation.

**Figure 1.**
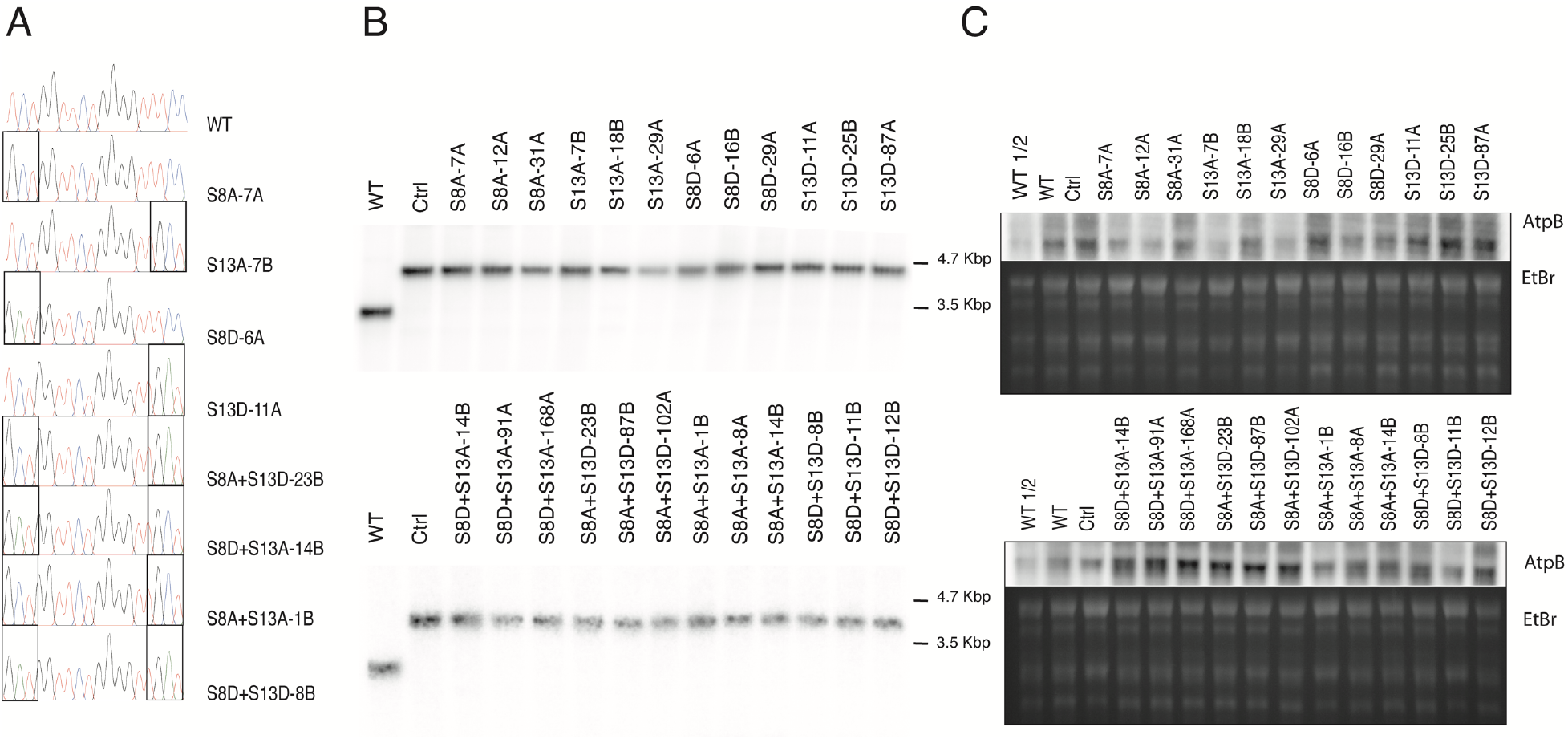
Molecular characterization of ATPβ mutants. (A) Sanger chromatograms from DNA of ~20 pooled seedlings resistant to spectinomycin; (B) Southern blot of DNA from A using a probe against *psaB;* (C) Northern blot of RNA from homoplasmic plants determined from A and B. Ctrl: *aadA* control line.

### Growth of Transplastomic atpB Mutants

As alterations in the activity of ATP synthase should impact the extent of *pmf* leading to changes in feedback regulation of electron transfer (Rott et al., 2011; Davis et al., 2016), we tested the hypothesis that modifications of S8 or S13 of ATPβ would alter plant growth due to altered feedback regulation in the light. Figure 2A shows representative growth of mutants harboring an alanine substitution. It appears that there is no defect in growth with one exception; S8A+S13D leads to a slower growing plant. As the S8A mutant and the S13A mutant have no growth impairment, this is likely due to the aspartic acid substitution for S13. Indeed, in Figure 2B we see representative growth for mutants harboring aspartic acid substitutions, and here we can clearly see that growth is only impaired when the aspartic acid replaces S13, with the single S13D mutation having the largest growth delay.

**Figure 2.**
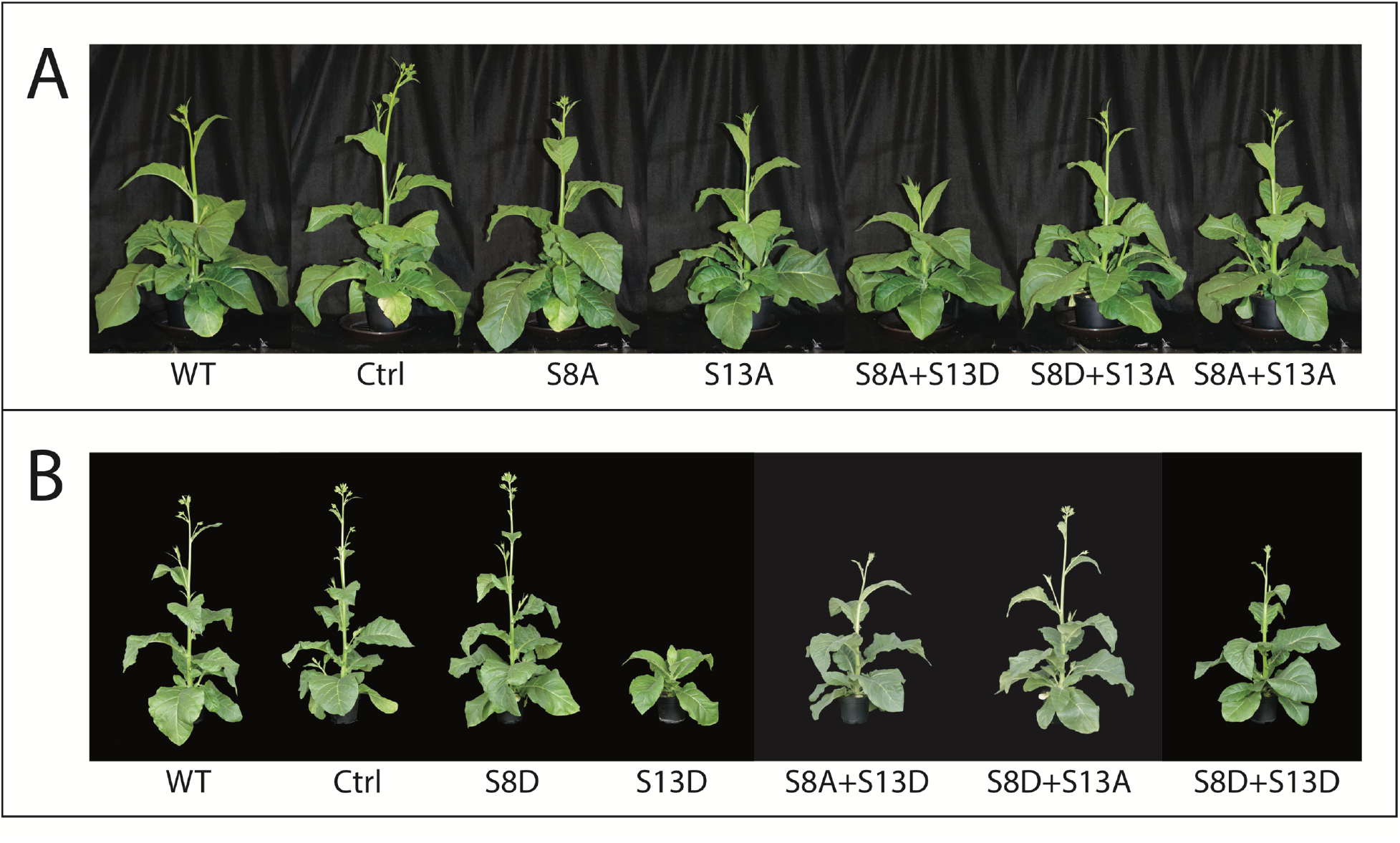
Representative growth of ATPβ mutants harboring alanine (A) or aspartic acid (B) substitutions on either S8 or S13. Lines used: (A) wild type (WT), Ctrl (*aadA* control line), S8A-7A, S13A-18B, S8A+S13D-102A, S8D+S13A-14B, S8A+S13A-8A; (B) WT, Ctrl, S8D-6A, S13D-87A, S8A+S13D-102A, S8D+S13A-168A, S8D+S13D-12B.

### Steady State Photosynthesis in the atpB Mutants

Since downregulation of photosynthesis due to altered ATP synthase activity was the most likely explanation for impaired growth in the mutants harboring the S13D mutation, we expected that we would see this downregulation reflected in photosynthetic parameters associated with *pmf* and PSII activity. To test this idea, we used chlorophyll *a* fluorescence to probe photosystem II (Baker, 2008), and the electrochromic shift (ECS) of thylakoid membrane embedded pigments in response to membrane potential (Bailleul et al., 2010) to probe the *pmf*. We hypothesized that the slower growing mutants (i.e., those with the S13D mutation) would have a decreased rate constant for proton efflux (*g*_H_^+^) which is a measure of ATP synthase activity, resulting in increased *pmf*, increased NPQ, and decreased photosynthetic efficiency (ϕ_II_). By taking the total extent of the dark to light changes of the ECS, we can calculate relative steady state *pmf* (ECS*_t_*, Figure 3A), and by fitting this decay to a first order exponential decay, we can calculate *g*_H_^+^ (Figure 3B) (Baker et al., 2007). After steady state illumination with ~400 μmol photons m^-2^ s^-1^, plants with the S13D mutation have increased relative total *pmf* (ECS*_t_*) (~150% increase for all S13D mutants; Figure 3B). This can be attributed to the decrease in *g*_H_^+^ seen for S13D plants (Figure 3A). The largest decrease in *g*_H_^+^ is seen in the single S13D mutant (~70% decrease), while the decrease in *g*_H_^+^ is comparable between the S8A+S13D and S8D+S13D mutants (~35% decrease). This decrease in *g*_H_^+^ also translates to an increase in feedback regulation in the form of NPQ (~110% increase for all S13D mutants; Figure 3C) and decreased photosynthetic efficiency (~70%, 60%, and 50% decrease for S13D, S8A+S13D, and S8D+S13D, respectively; Figure 3D), supporting our hypothesis that altered ATP synthase kinetics are the likely cause for the decreased growth of S13D mutants seen in Figure 2.

**Figure 3.**
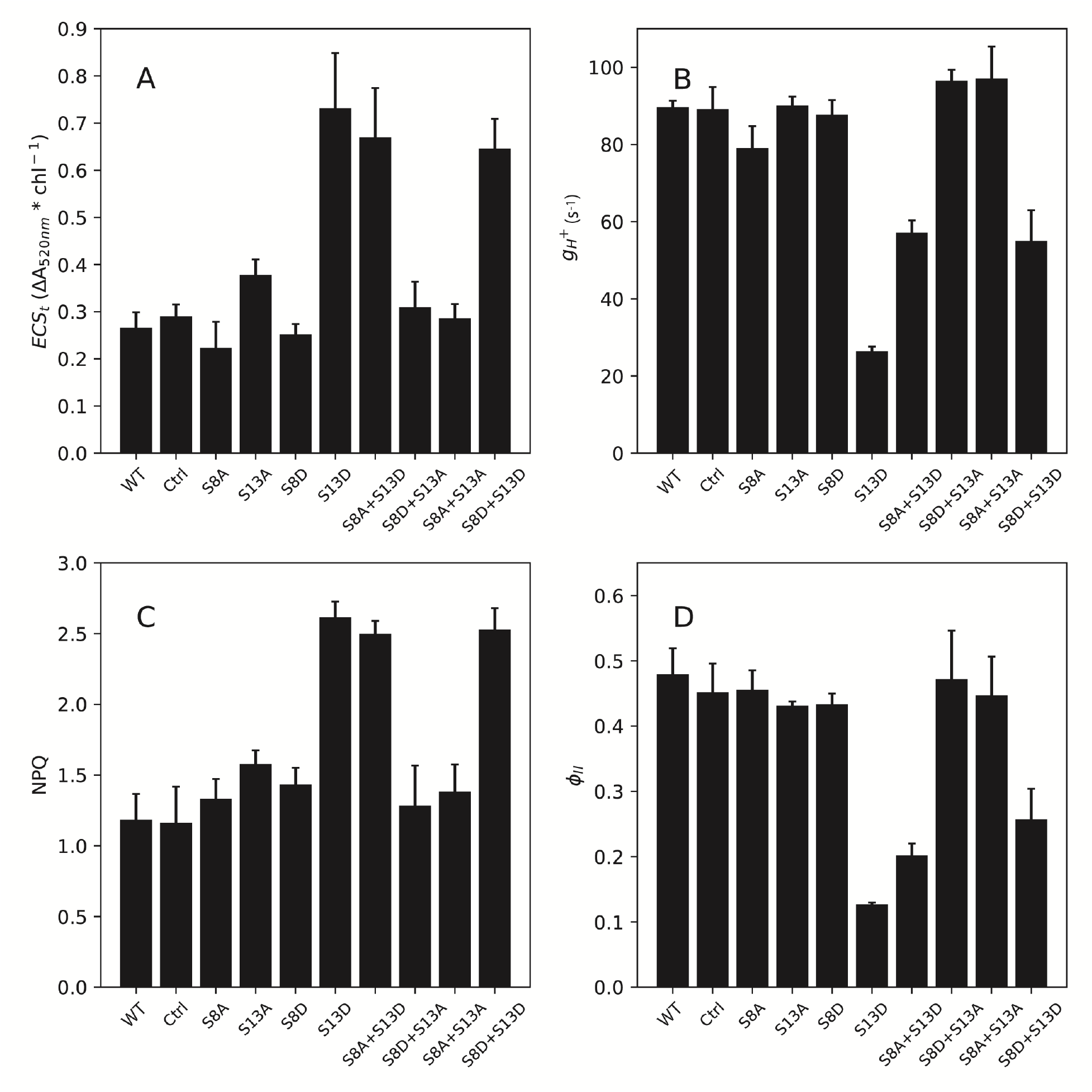
Steady-state photosynthetic parameters of ATPβ mutants in 400 μmoles photons m^-2^ s^-1^. (A) Total light to dark amplitude of the electrochromic shift at 520 nm; (B) Rate constant of proton efflux (*g*_H_^+^); (C) Nonphotochemical chlorophyll fluorescence quenching (NPQ); (D) quantum yield of photosystem II (ϕ_II_). Data represents mean +/- SD, n = 3. Lines used: *N. tabacum* Petit Havana (WT), Ctrl (*aadA* control line), S8A-31A, S13A-7B, S8D-29A, S13D-11A, S8A+S13D-23B, S8D+S13A-91A, S8A+S13A-8A, S8D+S13D-12B.

### ATP Synthase Protein Content in Transplastomic atpB Mutants

Due to the decrease in growth and changes in steady state photosynthetic parameters in the mutants harboring the S13D mutation, we tested two hypotheses: 1) the phosphorylation state of ATPβ has a regulatory function, as previously suggested (Reiland et al., 2009), and 2) the phosphorylation state alters ATP synthase content, which leads to a change in relative activity. In general, ATP synthase is thought to not be regulated by protein content. This is based on a previous study that determined at least 50% of ATP synthase content can be lost without a defect in steady state activity, as measured by *g*_H_^+^ (Rott et al., 2011). However, to rule out protein content changes as a cause for the decreased *g*_H_^+^ in the S13D plants, we quantified ATP synthase content of a CF_1_ subunit (ATPβ) and a CF_o_ subunit (ATPb, encoded by *atpF*) by tricine-SDS PAGE and western blot analyses (Figure 4).

**Figure 4.**
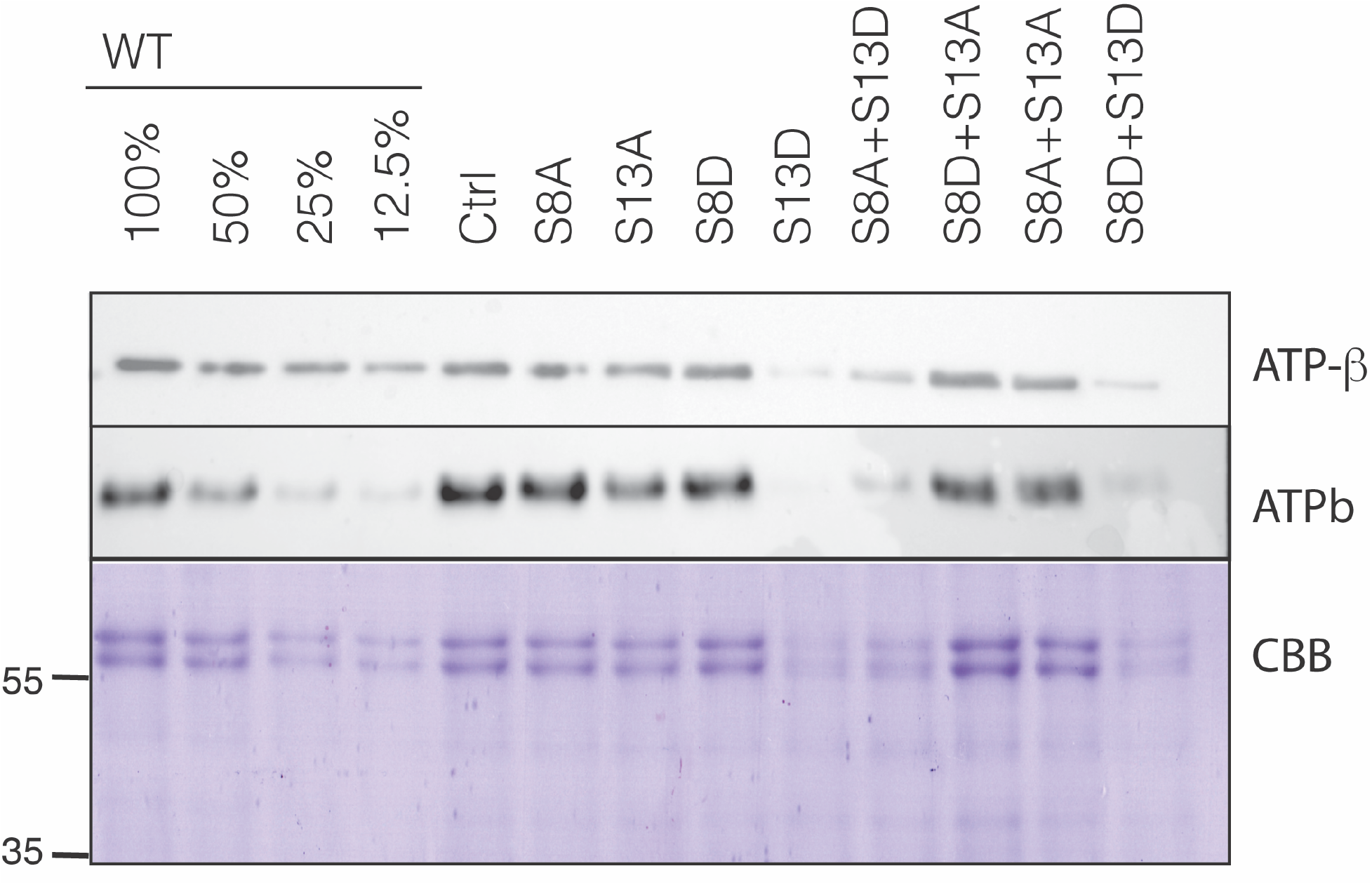
Immunoblot analysis of CF_o_ (ATPb) and CF_1_ (ATPβ) accumulation in isolated thylakoids of ATPβ mutants. ATPβ content is confirmed with Coommassie Brilliant Blue (CBB) staining of the α/β subunits (~60kDa). Lines used: *N. tabacum* Petit Havana (WT), Ctrl (*aadA* control line), S8A-31A, S13A-7B, S8D-29A, S13D-11A, S8A+S13D-23B, S8D+S13A-91A, S8A+S13A-8A, S8D+S13D-12B.

While the alanine mutations lead to no apparent change in the amount of ATPβ, this CF_1_ subunit accumulates to a lesser extent in mutants with the S13D substitution. ATPβ content in the S8A+S13D and S8D+S13D mutants is ~25% of WT and control plants, while the single S13D mutation leads to a reduction in ATPβ content to ~12.5% of wild-type levels. Since the ATPβ epitope is the mutated target, the western blot results were confirmed by Coomassie blue staining (Figure 4) where the α/β bands of the CF_1_ are seen clearly just above 55 kD. Further, the CF_o_ subunit (ATPb) accumulates similarly, with only the S13D mutations leading to a decrease in content (~25% of WT for the S8A+S13D and S8D+S13D, and ~12.5% for S13D). Considering these quantifications, we were therefore unable to rule out that loss of ATP synthase in the S13D mutants is the cause of the altered steady state *g*_H_^+^. It also appears likely that the stronger phenotype in the single mutation of S13D is due to this mutation leading to lower protein accumulation than the S8A+S13D and S8D+S13D mutants.

### ATP Synthase Dark Inactivation

The phosphorylation of ATPβ on residues S8 or S13 has previously been reported at the end of a long dark period (Reiland et al., 2009), which led the authors of that study to propose that phosphorylation is involved in the light to dark regulation of ATP synthase to prevent ATP hydrolysis in the dark. We set out to test this hypothesis by determining the inactivation kinetics of ATP synthase from light to dark. To this end, we measured several parameters involving ATP synthase activity that may change from light to dark. The best-known change in ATP synthase during a light to dark transition is the oxidation of the regulatory thiols on the g subunit. This oxidation leads to a decrease in the rate constant of the ECS decay after a weak flash in the dark (Kramer et al., 1990; Wu et al., 2007). Similarly, if phosphorylation plays a role in light to dark inactivation of ATP synthase, we might expect a decrease in the rate constant of this decay at earlier time points in the phosphomimic lines, and later time points in the alanine substitution lines. We thus applied a weak actinic flash to light-adapted leaves in increasing time increments after the constant illumination had been turned off and calculated the lifetime (τ_ECS_) of the decay from a first order exponential fit.

ATP synthase is not only redox regulated, but also requires *pmf* for activation (Kramer and Crofts, 1989). In prolonged dark, the threshold of *pmf* for activation (*pmf*_t_) increases as a function of time (Kramer and Crofts, 1989). It is not yet known what factors lead to an increase in *pmf*_t_ in the dark, but phosphorylation has not been ruled out. To test the hypothesis that the phosphorylation state of S8 and S13 on ATPβ increases the *pmf*_t_ in the dark, we gave a multiple turnover actinic flash and monitored the ECS decay kinetics in the leaves of our set of phosphorylation mutants. After the flash, the first derivative of the fast phase has a linear relationship with the amplitude of the flash induced ΔA_520 nm_, and the slope of this relationship is proportional to ATP synthase activity (*g*_H_^+^_d_), and the x-intercept is proportional to *pmf*_t_ (Kramer and Crofts, 1989). We grouped all lines according to substitution (i.e., A or D) and position (i.e., S8 or S13) to see which site/substitution resulted in an altered dark adaptation phenotype.

Figures 5 and 6 show dark inactivation parameters for all mutants harboring the S8A and S8D substitutions, respectively. We see that τ_ECS_ increases as expected, with the wild type and the control reaching ~2/3 maximum between 8-10 min (Figures 5 and 6, A). This oxidation state is maintained until prolonged darkness, where in the middle of the night (overnight, ON) the τ_ECS_, and thus g-oxidation reaches maximum. There is no clear agreement in τ_ECS_ deviation from wild-type and control kinetics for lines with S8A or S8D. In the wild type and control plants, *g*_H_^+^_d_ (Figures 5 and 6, B) is constant at all time points in the dark (ON). As seen with τ_ECS_, there is no clear deviation from wild-type and control kinetics caused by the S8A or S8D substitutions. Finally, *pmf*_t_ (Figures 5 and 6, C) increases with similar kinetics of τ_ECS_ in WT and control plants, with no clear differences in kinetics caused by either S8 substitution.

**Figure 5.**
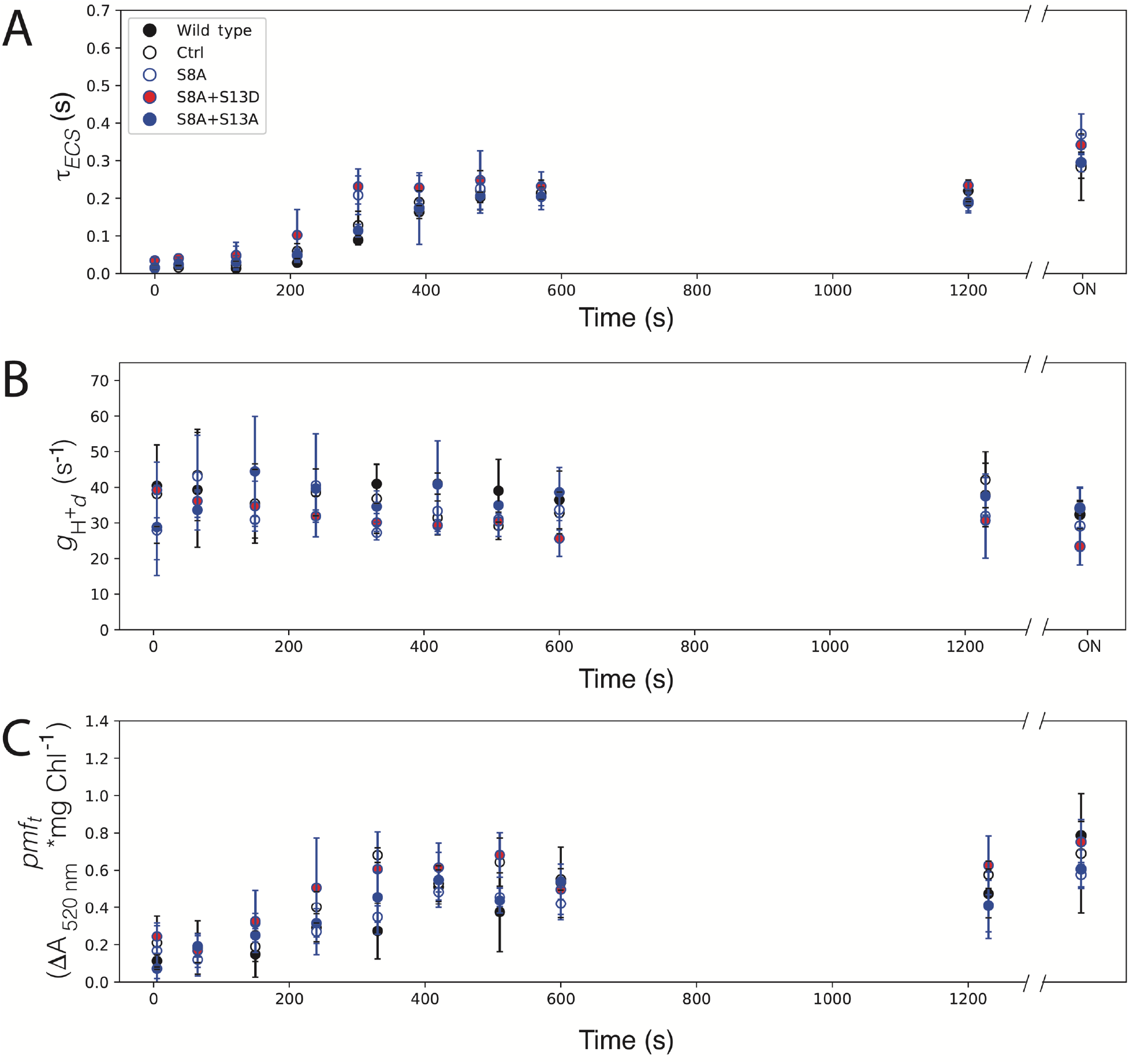
Kinetics of ATPβ -S8A mutants in the dark after illumination. (A) Lifetime of ECS decay (τ_ECS_) after a weak actinic flash, reporting the oxidation state of ATPγ. (B) *g*_H_^+^_d_. (C) *pmf*_t_. Data represents mean +/- SD, n = 3. Lines used: *N. tabacum* Petit Havana (Wild type), Ctrl (*aadA* control line), S8A-31A, S8A+S13D-23B, S8A+S13A-8A.

**Figure 6.**
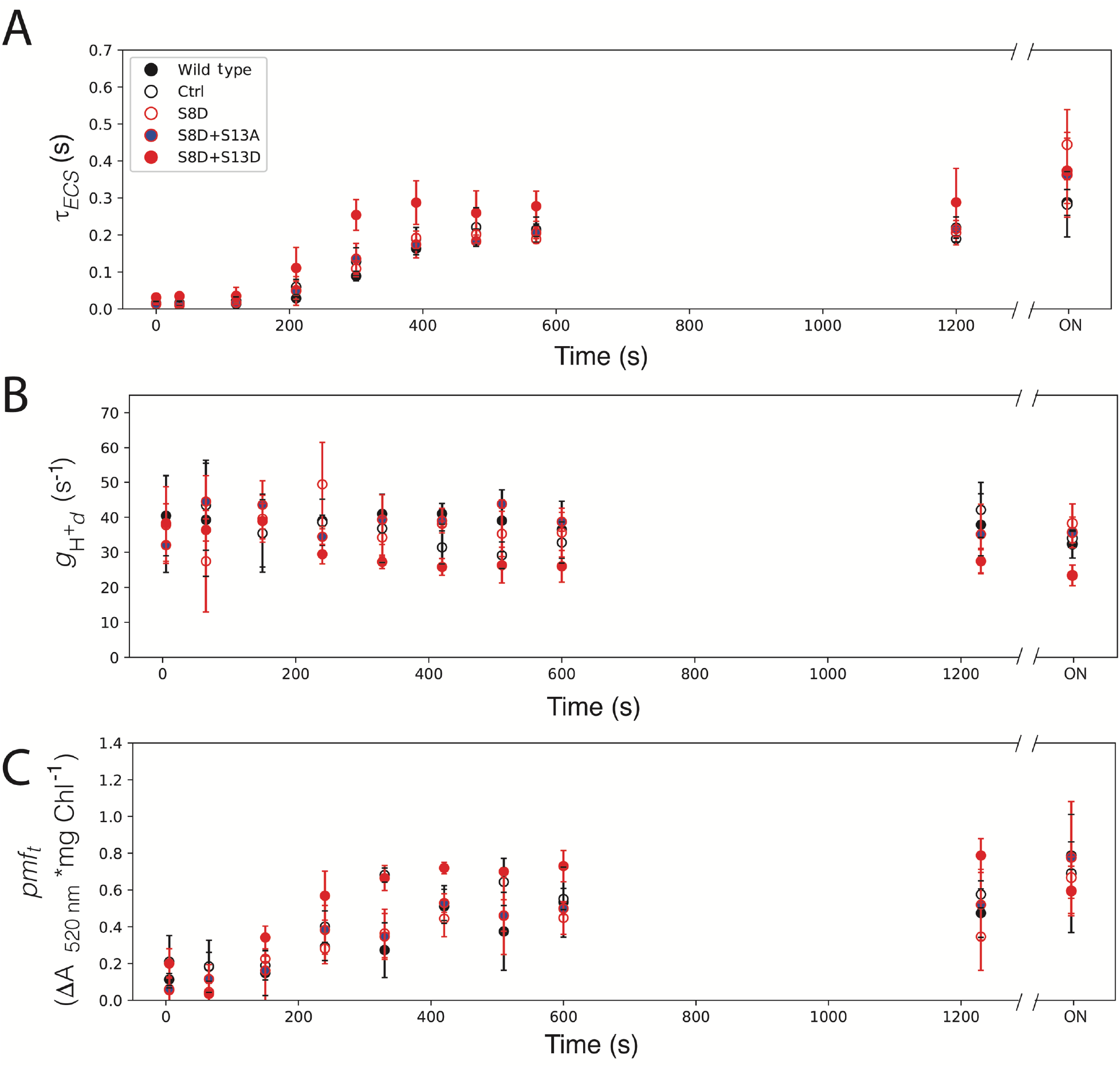
Kinetics of ATPβ -S8D mutants in the dark after illumination. (A) Lifetime of ECS decay (τ_ECS_) after a weak actinic flash, reporting the oxidation state of ATPγ. (B) *g*_H_^+^_d_. (C) *pmf*_t_. Data represents mean +/- SD, n = 3. Lines used: *N. tabacum* Petit Havana (Wild type), Ctrl (*aadA* control line), S8D-29A, S8D+S13A-91A, S8D+S13D-12B.

Figures 7 shows dark inactivation parameters for all mutants harboring the S13A substitutions. The S13A substitution results in all plants matching WT and control τ*_ECS_* (Figure 7A), *g*_H_^+^_d_, (Figure 7B), and *pmf*_t_ kinetics (Figure 7C). This suggests that elimination of this phosphorylation site has no impact on the dark inactivation of ATP synthase.

**Figure 7.**
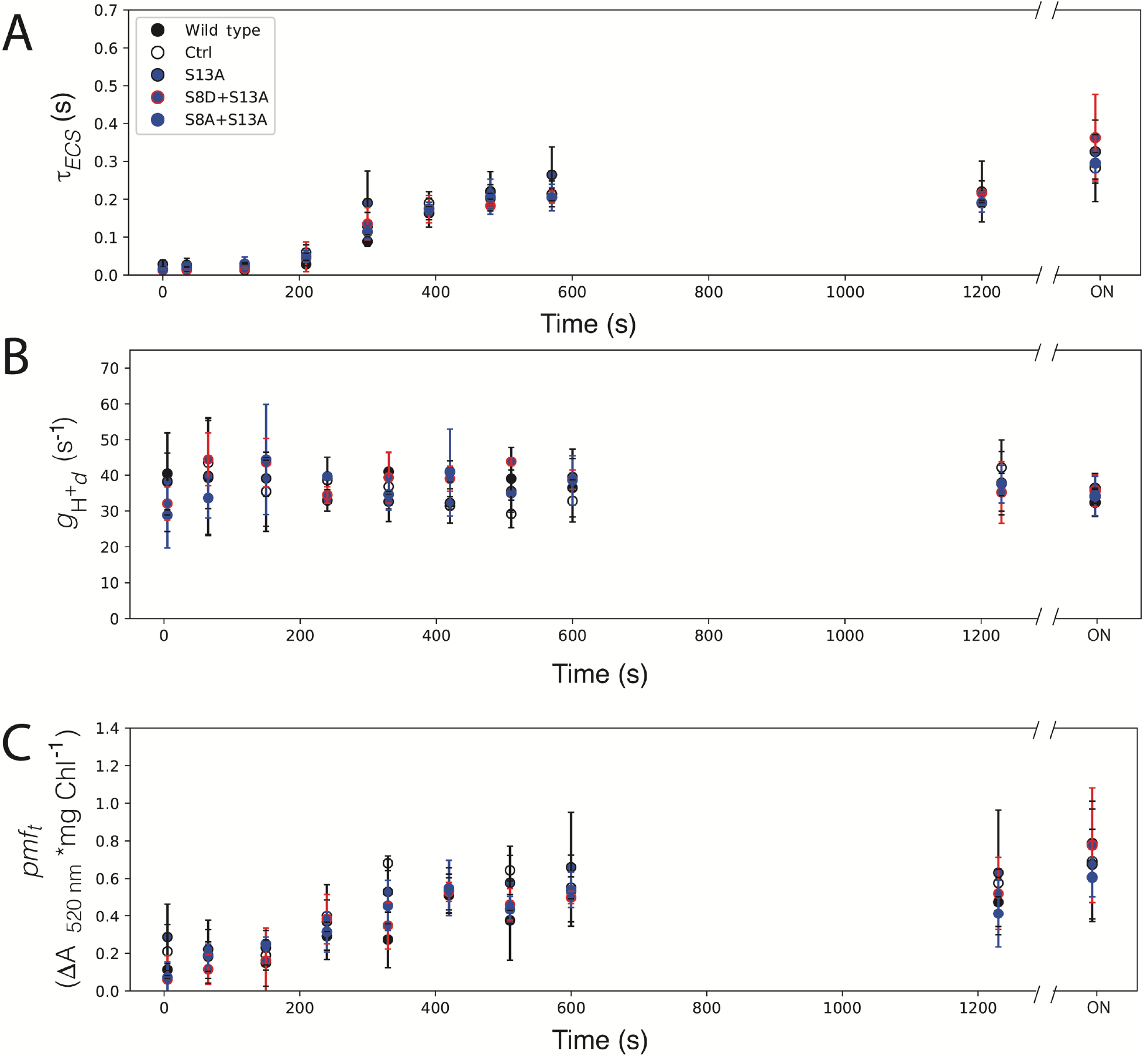
Kinetics of ATPβ-S13A mutants in the dark after illumination. (A) Lifetime of ECS decay (τ_ECS_) after a weak actinic flash, reporting the oxidation state of ATPg. (B) *g_H_^+^d*. (C) *pmf*_t_. Data represents mean +/- SD, n = 3. Lines used: *N. tabacum* Petit Havana (Wild type), Ctrl (*aadA* control), S13A-7B, S8D+S13A-91A, S8A+S13A-8A.

However, we see that when the S13D mutation is present, the prolonged intermediate oxidation state of ATPg is reached within 5 minutes of the switch from light to dark, compared to the ~10 min in the wild type and control plants. This finding indicates that g-oxidation is accelerated in the S13D plants (Figure 8A). There seems to be no clear trend to the final extent of g-oxidation (τ_ECS_ at time = ON) for either mutation or site, thus the variation is likely not reflective of anything related to the mutations or protein content. ATP synthase dark activity, *g*_H_^+^_d_, is lower in the S13D mutants than in the wild type and the *aadA* control lines (Figure 8B), with the strongest decrease from wild-type activity seen in the S13D single mutant. While in the WT, maximum *pmf*_t_ was reached after ~400 s in the dark and did not appear to increase to any significant extent until the ON timepoint, plants harboring the S13D mutation appear to have a faster increase in *pmf*_t_, reaching maximum *pmf*_t_ after around 5 min (Figure 8C). However, the ON differences between the wild type and control plants compared to the S13D mutants are unremarkable.

**Figure 8.**
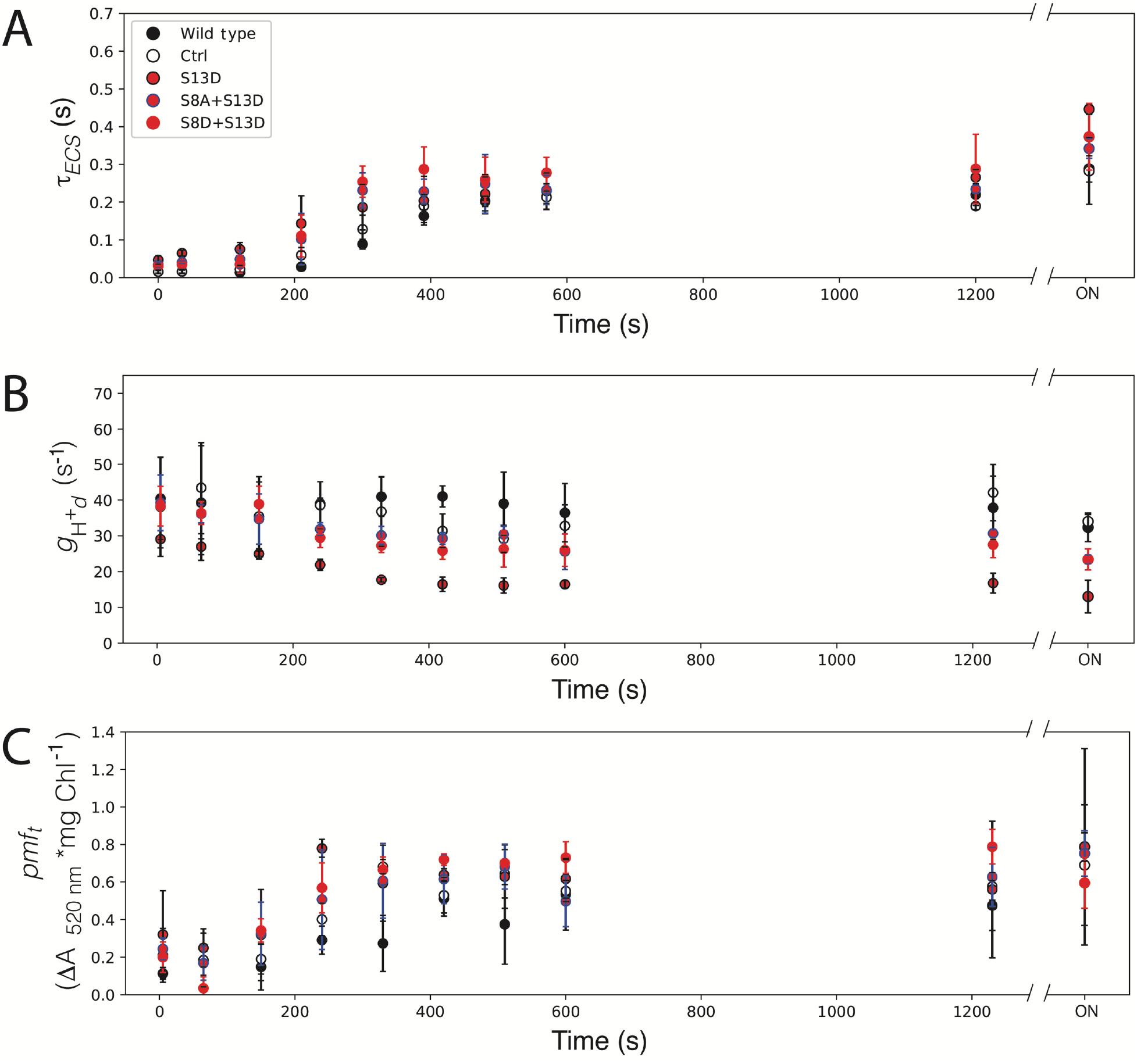
Kinetics of ATPβ--S13D mutants in the dark after illumination(A) Lifetime of ECS decay (τ_ECS_) after a weak actinic flash, reporting the oxidation state of ATPγ. (B) *g*_H_^+^_d_. (C) *pmf*_t_. Data represents mean +/- SD, n = 3. Lines used: *N. tabacum* Petit Havana (Wild type), Ctrl (*aadA* control), S13D-11A, S8A+S13D-23B, S8D+S13D-12B.

Since the only defect in ATP synthase inactivation kinetics was seen in plants with decreased ATP synthase content, we hypothesized that the altered kinetics were due to a decrease in protein content, and not regulation. We tested this hypothesis by performing similar experiments with the *ATPC1* antisense plants described in (Rott et al., 2011). These plants accumulate ~12% of ATP synthase, comparable in protein content with our most strongly defective S13 phosphomimic mutation. In these *ATPC1* antisense plants, we see a faster increase in τ_ECS_ (Fig 9A) in the dark, indicating that the decrease in ATP synthase content can indeed explain the increased rate of ATP-g oxidation in our S13 phosphomimic lines. Additionally, a decrease in *g*_H_^+^_d_ was seen in the ATPC1 knock-downs (Figure 9B), qualitatively mimicking the phenotype seen in the single S13D ATPβ mutant (Figure 8B). The *ATPC1* antisense lines also showed an earlier increase in *pmf*_t_ in the dark (Figure 9C). These data, taken together with the lack of a detectable phenotype of the S13A plants (Figure 7), suggest that the phenotypes seen for the S13D plants in the dark are not a result of the mimicked phosphorylation state, but instead, are due to the decreased ATP synthase content that results from the S13D mutation.

**Figure 9.**
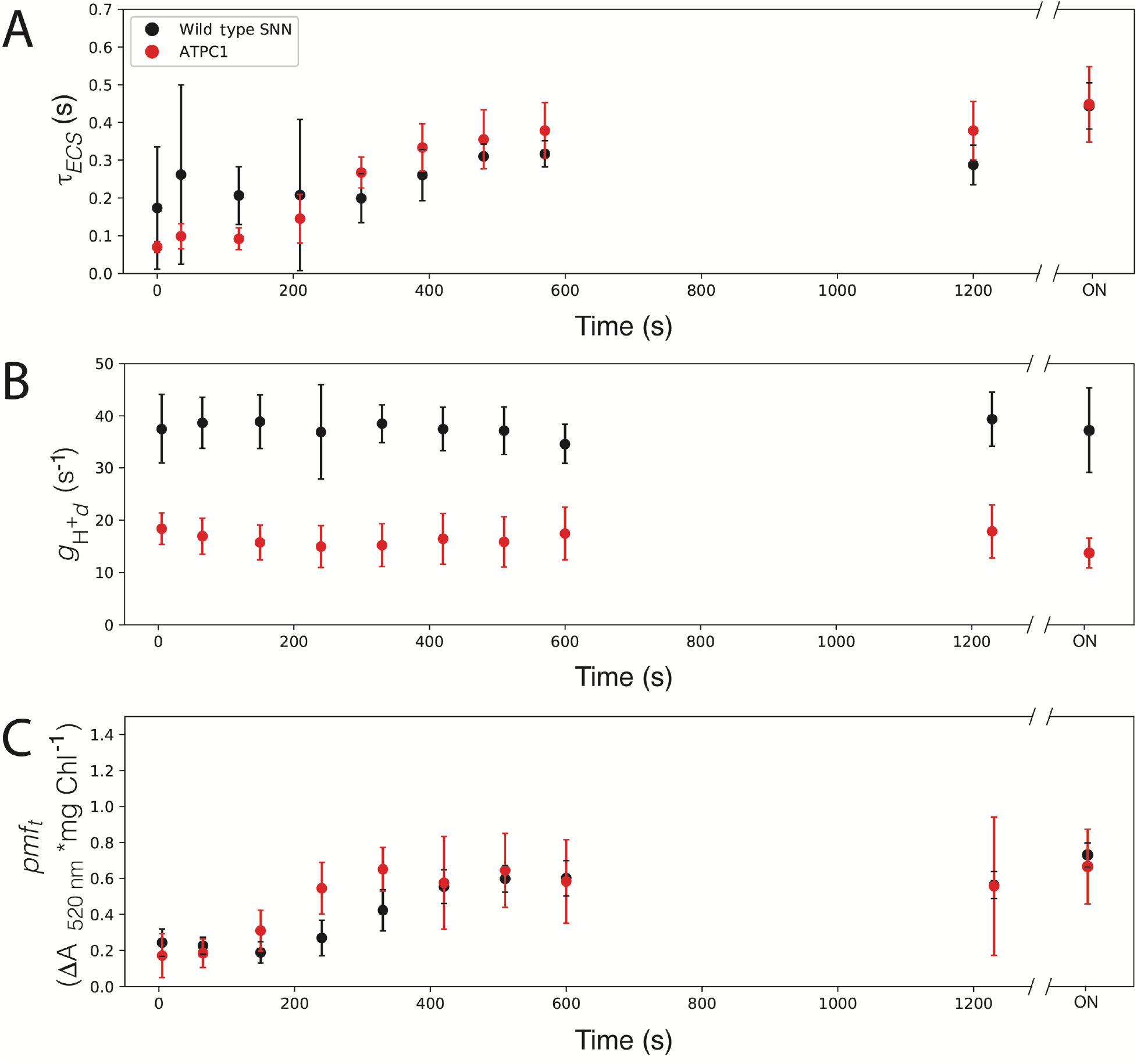
Kinetics of *ATPC1* knock-down mutants in the dark after illumination. (A) Lifetime of ECS decay (τ_ECS_) after a weak actinic flash, reporting the oxidation state of ATPγ. (B) *g*_H_^+^_d_. (C) *pmf*_t_. Data represents mean +/- SD, n = 3. Wild type SNN: *N. tabacum* cultivar SNN.

### ATP Synthase Content in Response to Drought

In the absence of any significant change to ATP synthase dark inactivation in our non-phosphorylateable mutants, we considered conditions in which ATP synthase activity or content is altered to test the hypothesis that phosphorylation is involved in modulating activity under stress conditions. Known conditions under which ATP synthase activity is altered include response to drought (Kohzuma et al., 2009), low CO_2_ (Kanazawa and Kramer, 2002), and phosphate limitation (Takizawa et al., 2008). Under drought stress, ATP synthase content and activity is decreased (Kohzuma et al., 2009), and ATPβ has been shown to be phosphorylated by CK-II in vitro (Kanekatsu et al., 1998), a protein kinase proposed to be involved in drought stress responses (Vilela et al., 2015). These observations led us to hypothesize that the phosphorylation of ATPβ could be a signal for the decrease in ATP synthase content in response to drought. To test this idea, we drought-stressed our mutants harboring the S➔A mutations and measured ATPβ content by western blot analysis. Figure 10 shows that all mutants have a decrease in ATPβ content after drought stress, regardless of the mutation, indicating that phosphorylation of the S8 or S13 residues is unlikely to be the signal for decreased ATP synthase content in response to drought.

**Figure 10.**
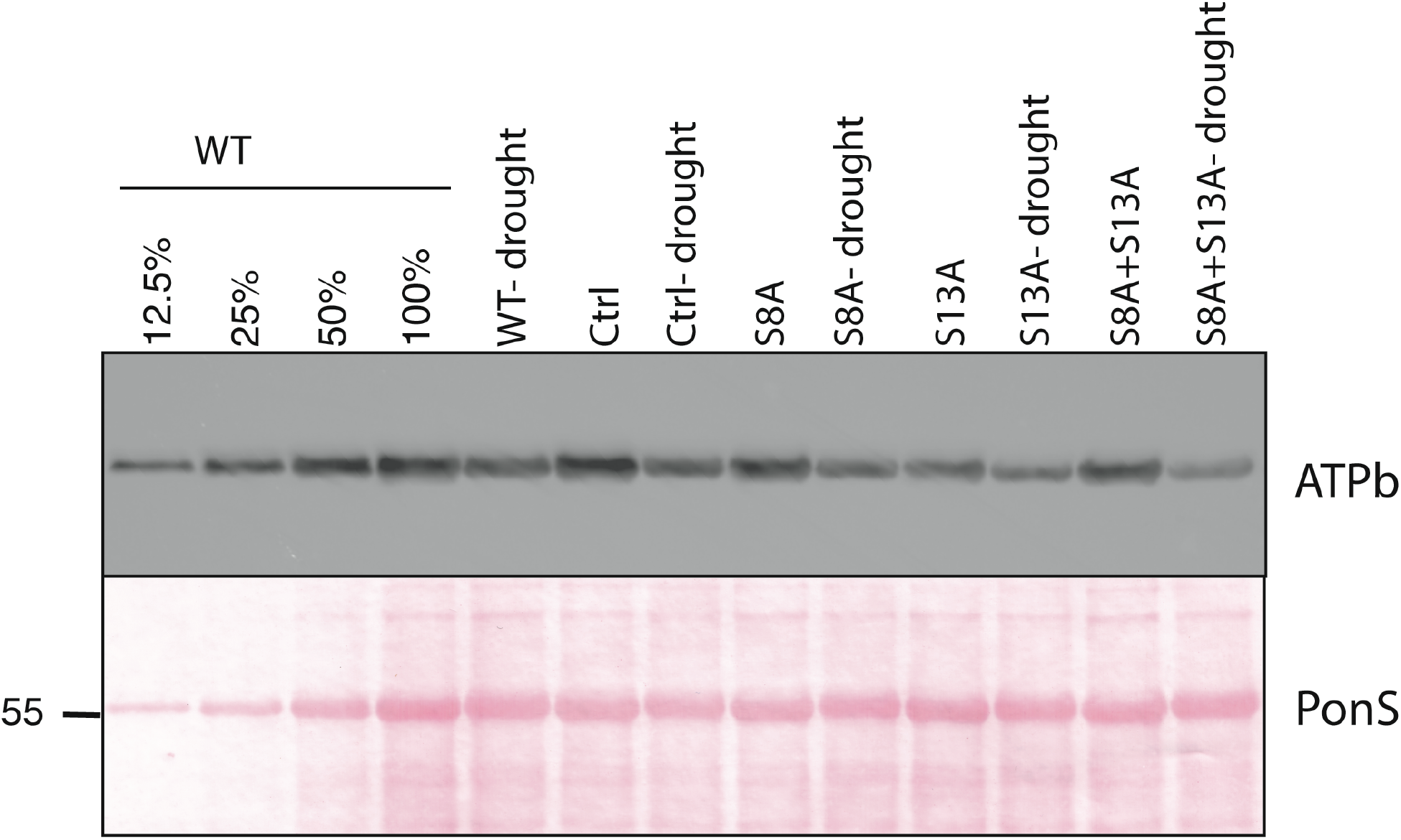
Immunoblot analysis of CF_o_ (ATPb) protein accumulation in total leaf protein samples from well-watered and drought stressed plants. Lines used: *N. tabacum* Petit Havana (WT), Ctrl (aadA control line), S8A-12A, S13A-29A, S8A+S13A-14B. PonS: Ponceau S staining of the membrane after blotting.

### Kinetics of ATPb Turnover in ATPβ Mutants

To understand the decreased accumulation of ATP synthase subunits in our S13 mutants, we generated two new hypotheses. 1) The S13D substitution leads to an increase in the degradation rates of ATP synthase, and 2) the S13D substitution leads to a decrease in the rate of ATP synthase synthesis. To test the hypothesis that the mutations alter turnover rate of ATP synthase, we infiltrated leaf discs with lincomycin and sampled at 24, 48, 96, and 144 hours after infiltration. Total protein samples from our time course were then blotted and hybridized to an antibody against ATPb (Figure 11). We hypothesized that we would see a higher rate of ATPb degradation in the S13D mutants and a slower rate of ATPb degradation in the S13A mutants, if phosphorylation was signaling for protein turnover. Figure 11 shows the ATPb degradation kinetics for our ATBβ mutants that show wild-type ATP synthase accumulation. Degradation kinetics appear to be wild type-like for the control plants and the S8A+S13A mutant (Figure 11A), the S8A and S13A mutants (Figure 11B), and the S8D and S8D+S13A mutants (Figure 11C). To see the degradation kinetics of ATPb in our low accumulating ATPβ mutants (i.e., mutants harboring the S13D mutation), we increased the protein loading to levels that would give us near wild-type signals for ATPb (400% of WT for S8A+S13D and S8D+S13D, and 800% of WT for S13D; Figure 11D and E, respectively). On a total protein basis, ATPb does not appear to decrease at a substantially different rate for the S13D mutants. We, therefore, rejected the hypothesis that changes in ATP synthase content in our phosphomimic lines is due to increased protein degradation rates. Instead, it seems likely that the lower ATP synthase content in the S13D mutants is due to some step during synthesis and/or assembly of the complex, especially considering that there appear to be no defects in mRNA expression or stability (Figure 1C).

**Figure 11.**
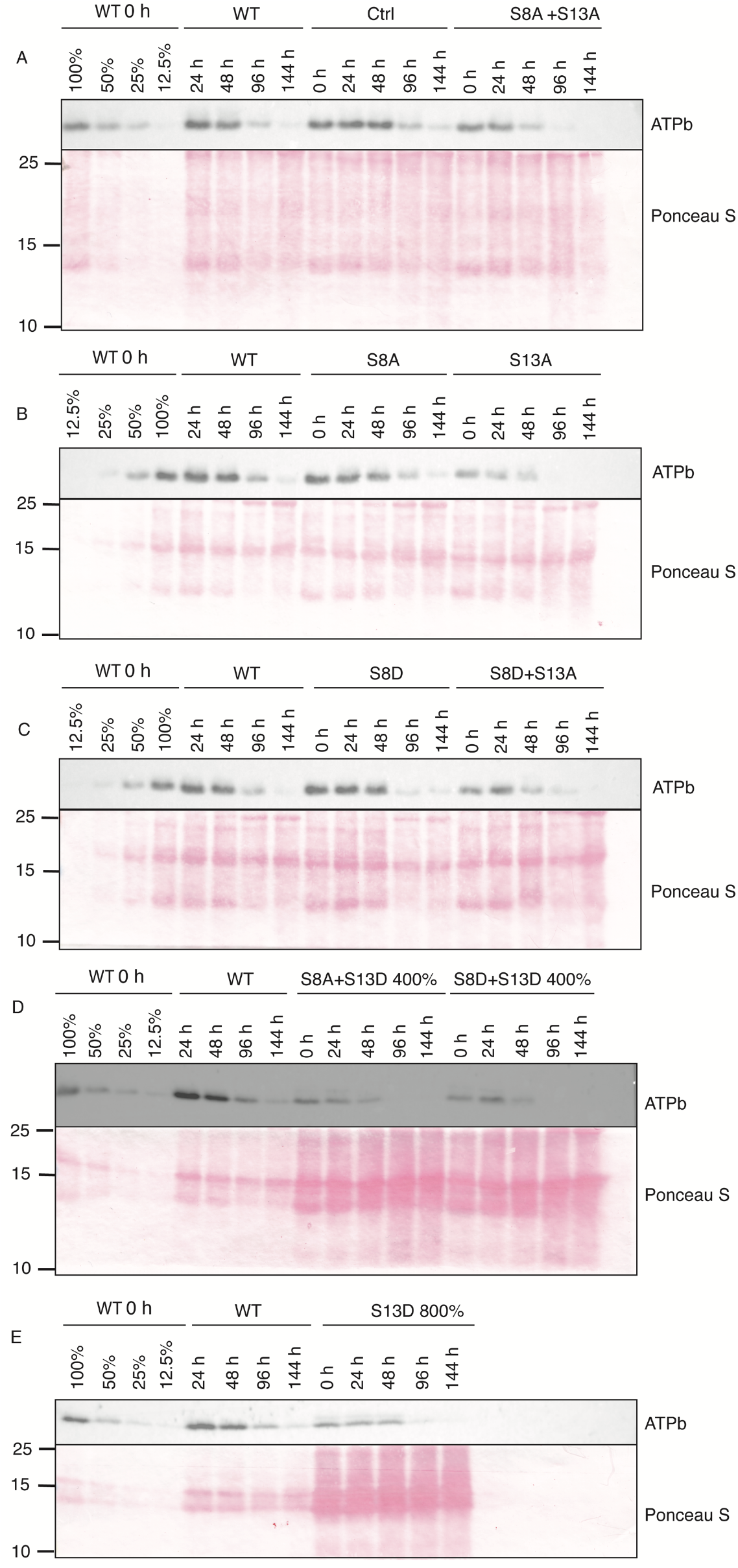
Time course of ATPb content after infiltration of leaf discs with 3 mM lincomycin. Lines used: *N. tabacum* Petit Havana (WT), Ctrl (*aadA* control line), S8A-7A, S13A-29A, S8D-16B, S8D+S13A-91A, S8A+S13A-14B, S13D-11A, S8A+S13D-102A, S8D+S13D-8B.

## Discussion

To assess the role of N-terminal phosphorylation of ATPβ in photosynthetic regulation, we used a set of mutants locked in specific phosphorylation states. We saw defects in ATP synthase in the light and altered inactivation kinetics in the dark associated with decreased protein accumulation in all mutants harboring the S13D mutation. When we tested the hypothesis that the phosphorylation state of ATPβ serines 8 and 13 contributed to the dark inactivation of ATP synthase as proposed in (Reiland et al., 2009), we only saw changes in S13D plants (Figure 8). These plants had faster ATPg oxidation and appearance of increased *pmf*_t_, and in the case of the single S13D mutation, lower ATP synthase activity in the dark (*g*_H_^+^_d_). We demonstrated that this is due to loss of ATP synthase complex, and not dynamic regulation of the complex, by comparing the inactivation kinetics of an ATP synthase knock-down mutant (Figure 9, (Rott et al., 2011)), which yielded qualitatively similar results. Therefore, we rejected the hypothesis that phosphorylation contributes to inactivation in the dark. Instead, the main light/dark switch in the chloroplast ATP synthase is likely oxidation of ATPg, as has been demonstrated elsewhere (Kohzuma et al., 2012). In fact, it seems that the ATPg oxidation state is likely the regulator of the *pmf* threshold of activation (*pmf*_t_) for ATP synthase, as both of these are altered on the same timescale when ATP synthase content is decreased. In general, the amount of ATP synthase in the chloroplast should not alter the Em of ATPg. Instead, the faster oxidation of ATPg could be explained by a decrease in electron transfer that might be expected from the decrease in ϕ_II_ (Figure 3) we see in the S13D plants. A decrease in the bulk flow of electrons may lower the electron buffering capacity of the stroma in the dark, leading to quicker oxidation of ATPg and likely other redox-regulated proteins in the chloroplast.

Due to the decrease in ATP synthase content in our ATPβ S13D mutants (Figure 4), we hypothesized that phosphorylation was a signal for protein degradation in the dark. However, in the presence of lincomycin, there was no substantial differences in the apparent degradation rate of the CF_o_ subunit ATPb in any of our mutants (Figure 11). In addition, our mutants accumulate wild type-like transcript levels (Figure 1C), suggesting that the defect in protein accumulation is either due to reduced translation or impaired assembly. It seems possible that degradation involves a chloroplast-encoded factor that would be inhibited by lincomycin, such as the P1 subunit of the Clp protease (Moreno et al., 2018), but ATP synthase is currently not thought to be a target of Clp. Therefore, with the information we have on hand, we must reject our hypothesis that the phosphorylation state of ATPβ-S13 is a signal for degradation.

As the rate of translation is unlikely to be altered due to a change in a single internal codon, the remaining point for a defect in complex accumulation is assembly. There is no evidence suggesting misassembly of ATP synthase in the S13D mutants. CF_o_ and CF_1_ subunits appear to be lost stoichiometrically (Figure 4), whereas misassembly would likely lead to a leaky enzyme and result in slowed ATPg oxidation kinetics in Figure 8A (as discussed in (Strand et al., 2017)). Thus, the defect to accumulation appears to be kinetic only. As of writing, no X-ray crystal or cryo-EM structure has been resolved containing the S8 or S13 residues of CF_1_-β, suggesting that these amino acids lie within a disordered region of the CF_1_. Given this lack of structural data, it is difficult to further hypothesize what, if any, defect the phosphomimics are causing in ATP synthase assembly.

In this manuscript, we ruled out multiple hypotheses for the role of ATPβ N-terminal phosphorylation in ATP synthase regulation, specifically the hypothesis of light-dark regulation and protein turnover. There are other possible roles for this phosphorylation in ATP synthase regulation that have yet to be tested. As ATP synthase is proposed to have multiple regulatory phosphorylation sites, identification of phosphorylation of specific ATP synthase residues under conditions in which activity is known to be altered (e.g., low CO_2_ or P_1_) may be a more productive first approach. It has been proposed that CK-II is a kinase for ATPβ (Kanekatsu et al., 1998; Reiland et al., 2009), and phosphorylation still may serve some yet unknown function in the dark. ATPβ forms part of the catalytic domain mediating ATP synthesis and hydrolysis. If we hypothesize that phosphorylation of these residues is functional and not artifactual or promiscuous, then the question becomes ‘what occurs at night that would require regulation of the catalytic CF_1_ domain?’. It has been proposed that ATP synthase needs to be inactivated so that dark ATP hydrolysis does not occur (Kohzuma et al., 2012). However, since ATP hydrolysis in the dark is linked to *pmf* formation, and we do not see any *pmf* linked changes in activity that cannot be attributed to protein content, the activity that is being regulated would likely need to be independent of *pmf*. One possible function for phosphorylation in the dark could be to shift the K_M_ of substrate to allow for changes in the activation kinetics of ATP synthase in the morning, and it may be that a more dynamic light regime would reveal an impact on plant growth in those mutants impaired in phosphorylation.

An interesting aspect of ATP synthase is that it is regulated primarily post-translationally, and the rate constant of proton efflux has been shown to be unresponsive to decreases in ATP synthase content as low as 50%, suggesting that there is a pool of inactive ATP synthase in the thylakoid (Rott et al., 2011). In that report, the relationship between activity and content remained unknown between 12.5 and 50%. Therefore, the extent of the inactive fraction of the ATP synthase pool was assumed to be 50% due to the linear relationship of activity and protein content between 50% and ~12.5%. Here we contribute another point to the curve, 25% ATP synthase content, shown for our ATPβ mutants S8A+S13D and S8D+S13D (Figure 4). We see that a 75% decrease in ATP synthase content leads to a ~40% decrease in activity, while the 87.5% decrease in content in our S13D mutant leads to a ~75-80% decrease in activity (Figures 4), similar to the decrease in activity seen in (Rott et al., 2011) with similar ATP synthase content. These data suggest no regulatory role of N-terminal ATPβ phosphorylation in photophosphorylation, and the simplest explanation for decreased *g*_H_^+^ is the decrease in ATP synthase content. Since we show that ATP synthase activity decreases at 25% content, we suggest an inactive fraction of between 50% and 75%. However, all else being equal, if we assume that the inactive fraction is 50%, there is a roughly linear relationship between rate constant and protein content when the inactive pool is eliminated. However, if the inactive pool is >50%, the relationship appears more exponential like, which would suggest that the remaining ATP synthase in these mutants are less active than their wild-type counterparts, an unlikely scenario.

In summary, our work reported here (i) casts considerable doubt on the proposed crucial role of the phosphorylation of amino acid residues Ser8 and Ser13 in the regulation of ATP synthase activity, and (ii) suggests a function of amino acid 13 in CF_1_ assembly.

## Methods

### Plant material and growth conditions

Leaf tissue for DNA analyses came from plants grown on 0.5 MS medium (6.8 g/L phytoagar, 3% sucrose) at 50 μmol photons m^-2^ s^-1^. For RNA and protein analyses, plants were grown in 200 μmol photons m^-2^ s^-1^ with a 16:8 light:dark photoperiod. RNA was extracted from the youngest leaf of plants at 3 weeks after sowing. Spectroscopic analysis and protein extraction was performed on fully expanded leaves (17-20 cm in length) at ~5 weeks after sowing. For growth comparison, seeds were germinated in 200 μmol photons m^-2^ s^-1^ with a 16:8 light:dark photoperiod for approximately 3 weeks, then transferred to larger pots and grown at 350 μmol photons m^-2^ s^-1^ for approximately 8 weeks. Photos were taken when control plants (transformed with the construct containing the *aadA* marker but no *atpB* mutation) and wildtype plants began producing flower buds. Due to space constraints, representative lines harboring the S-to-A mutations at either site and representative lines harboring the S-to-D mutations at either site, were grown in independent experiments.

### Construction of plastid transformation vectors and plastid transformation

A 2018 bp region of the *Nicotiana tabacum* Petit Havana plastid DNA containing the coding sequence for the N-termini of ATPβ and RbcL was amplified by PCR and inserted into cloning vector pMCS5 (MoBiTec) between the PmeI and PacI restriction sites (NEB) using Gibson assembly (Gibson, 2011) and primer pair P1 and P2 (see Supplementary Table 1). The resulting plasmid (pDDS026) was then used as template to introduce all mutations except S8D+S13A by site directed mutagenesis using mismatched primer pairs (Supplementary Table 1). After amplification, the PCR products were digested with the restriction enzyme DpnI (NEB) to eliminate the template plasmid. To obtain the S8D+S13A mutation, a Gibson assembly protocol was employed using primer pairs P1/P17, and P2/P18, with P17 and P18 incorporating the two desired point mutations (Supplementary Table 1). The resulting fragments were inserted into pMCS5 as described for pDDS026. In total, 8 combinations of mutations were generated (Supplementary Table 1) and transformed into *E. coli* strain TOP10. Ampicillin-resistant colonies were sequenced for the desired mutation(s) before being combined with plastid transformation vector pDK308 (Nt-EctWT, (Agrawal et al., 2020)), containing a chimeric *aadA* gene (Svab and Maliga, 1993) as selectable marker gene for chloroplast transformation, and used in biolistic co-transformation experiments as described in (Fuentes et al., 2012; Krech et al., 2013).

### Selection of transplastomic tobacco lines

After particle bombardement, primary transplastomic lines were selected as described in (Caroca et al., 2021). Plants that appeared heteroplasmic by PCR and Sanger sequencing underwent additional regeneration rounds until they appeared homoplasmic by sequencing. Putative homoplasmic plants were further analyzed by Southern blot for homoplasmic integration of the *aadA* cassette, and lines with a single band shift of 1.2 kb were transferred to soil for seed production. Homoplasmy was confirmed for integration of the *aadA* cassette by plating the resulting seeds on selection medium and Southern blot analysis of DNA from ~20 pooled seedlings. Lines homoplasmic for the *aadA* displayed a homogeneous population of spectinomycin-resistant progeny (Bock, 2001) and a single band at 5.7 kb in the Southern blot analysis. Homoplasmy for the point mutations was confirmed by PCR amplification and Sanger sequencing of the same DNA as used for Southern blot analysis. Lines were considered homoplasmic when they had a single peak in the sequence chromatograms.

### Isolation of nucleic acids and gel blot analyses

Leaf tissue for DNA and RNA analyses was frozen in liquid nitrogen and pulverized using a steel ball Retsch mill. DNA was extracted according to (Doyle and Doyle, 1987) for Sanger sequencing and Southern blotting. For Southern blot analyses, total genomic DNA was digested with BglII, separated by gel electrophoresis in 0.7% (w/v) agarose gels and blotted onto Hybond N nylon membranes (GE Healthcare). A ~500 bp probe was produced by PCR amplification (primer pair P19/P20) of part of the *psaB* gene, followed by [α-^32^P]dCTP-labeling by random priming (Multiprime DNA labeling kit; GE Healthcare).

Total RNA extraction and northern blot analysis was performed as described in (Rott et al., 2011) using primer pair 21/22 (Table 1) to generate a DNA probe against *atpB*.

### Thylakoid extraction

Crude chloroplasts were extracted from mature leaves (~15-20 cm in length) as described in (Strand et al., 2016) with the addition of 10 mM sodium ascorbate to all buffers. Chloroplasts were then subjected to an osmotic shock (10 mM HEPES, 5 mM MgCl_2_, 2.5 mM EDTA, 10 mM sodium ascorbate, pH 7.6) on ice for 10 min and thylakoids were pelleted at 3000 x g for 10 min at 4°C. The pellet was resuspended in the osmotic shock buffer and used for subsequent experiments.

### Protein sample preparation and Tricine-SDS PAGE

For western blots based on chlorophyll concentration, thylakoids of the desired chlorophyll concentration were centrifuged at 3000 x g for 10 min at 4°C and resuspended in a modified Laemmli sample buffer (100 mM Tris-HCl, 4% SDS, 8 M urea, 12% β-mercaptoethanol, and 20% glycerol). For blots based on total protein, leaf punches from mature leaves were ground in liquid nitrogen in a Retsch mill. Protein was then extracted in buffer (100 mM Tricine-KOH, 2 mM MgCl_2_, 10 mM NaCl, 1 mM EDTA, 0.2% Trition-X 100, pH 7.5) and assayed for total protein content (Protein Assay Kit I, Bio-Rad). The protein was then precipitated in 80% acetone at −20°C overnight, and further treated identically to the samples loaded based on chlorophyll concentration.

Protein samples were gel electrophoretically separated as described in (Schägger, 2006) through a 4% stacking and 10% resolving gel at 16°C. Proteins were transferred to a nitrocellulose membrane with 350 mA for 4 h, with the voltage not exceeding 20 V, using a semidry transfer apparatus and transfer buffer containing 48 mM Tris, 39 mM glycine, and 20% methanol. Membranes were stained with Ponceau S prior to blocking. Membranes were then rinsed in deionized water and incubated for 1 h in TBST containing 5% dry milk as a blocking agent. Antibodies (anti-AtpB for ATPβ; anti-AtpF for ATPb; Agrisera) used to probe the membranes were prepared according the manufacturer’s dilution recommendation, and incubated on the membrane overnight at 4°C. Chemiluminescense from ECL Prime Western Blotting Detection Reagents (GE Healthcare) was detected on the G:Box Chemi XT4 (Syngene).

### In vivo absorption and fluorescence spectroscopy

Measurements were performed on a home-built spectrophotometer described in (Hall et al., 2013). Steady state chlorophyll a fluorescence and absorbance measurements were performed as described in (Strand et al., 2017). Plants were dark adapted for 30 min prior to experimentation. Actinic illumination was 400 μmol photons m^-2^ s^-1^.

To measure ATP synthase inactivation kinetics, two flash-induced relaxation kinetics (FIRK) of the electrochromic shift at 520 nm were performed as described in (Kramer and Crofts, 1989). Plants were preilluminated with 400 μmol photons m^-2^ s^-1^ prior to experimentation. After 15 min of illumination, the light was turned off. First, a weak flash was given at 0, 35, 120, 210, 300, 390, 480, 570, and 1200 s post illumination, and the absorbance change from a green LED filtered with a 520 nm bandpass filter was monitored. The decay was fit to a first order exponential decay to determine the lifetime (τ*_ECS_*). Second, a sub-saturating flash was given at 5, 65, 150, 240, 330, 420, 510, 500, and 1230 s post illumination, and the absorbance change at 520 nm was monitored. The first derivative of the decay was plotted against the amplitude of the DA520 nm. The slope of the linear portion was taken as the rate constant of ATP synthase in the dark (*g*_H_^+^_d_), and the x-intercept (ΔA_520 nm_ amplitude) was taken as *pmf* threshold of activation (*pmf*_t_). For the overnight dark adaptation, plants were measured for all FIRK parameters at least 4 h after the growth chamber lights had turned off.

All amplitude-dependent ECS (ΔA_520 nm_) parameters were normalized to chlorophyll concentration of the leaf area measured.

## Abbreviations

CBB: Coommassie Brilliant Blue
CF_o_: membrane fraction of ATP synthase
CF_1_: soluble fraction of ATP synthase
*pmf*: protonmotive force
q_E_: exciton quenching
NPQ: nonphotochemical quenching
ΔpH: fraction of *pmf* stored as pH
*bf*: cytochrome *b_6_f* complex
Q_o_: quinol oxidation site
WT: wild type
PCR: polymerase chain reaction
ATP: adenosine triphosphate
DNA: deoxyribonucleic acid
RNA: ribonucleic acid
ECS: electrochromic shift
ECS*_t_*: total electrochromic shift
τ_ECS_: lifetime of the electrochromic shift
ϕ_II_: quantum yield of photosystem II
*g*_H_^+^: transthylakoid proton conductivity
PSI: photosystem I
PSII: photosystem II
*g*_H_^+^_d_: transthylakoid proton conductivity in the dark
*pmf*_t_: protonmotive force threshold of activation
ON: overnight
CO_2_: carbon dioxide
NADPH: nicotinamide adenine dinucleotide phosphate

## Acknowledgements

We thank Margit Röβner and Peggy Gabor (MPI-MP) for help with tissue culture and transformation and the MPI-MP GreenTeam for plant cultivation. We thank Stephanie Seeger (MPI-MP) for help with RNA extraction and northern blotting. We thank Drs. Ute Armbruster and David Kramer for helpful discussions. This work was supported by a grant from the Deutsche Forschungsgemeinschaft to R.B. (BO 1482/17-2; FOR 2092) and by the Max Planck Society.

## Author contributions

**Deserah D. Strand:** Conceptualization, Methodology, Writing **Daniel Karcher:** Methodology **Stephanie Ruf**: Methodology **Anne Schadach:** Methodology **Mark A. Schöttler:** Conceptualization **Omar A. Sandoval Ibanez:** Conceptualization, Methodology **Ralph Bock:** Conceptualization, Methodology, Writing

